# Medial prefrontal activity at encoding determines enhanced recognition of threatening faces after 1.5 years

**DOI:** 10.1101/2020.08.25.266353

**Authors:** Xiqin Liu, Xinqi Zhou, Yixu Zeng, Jialin Li, Weihua Zhao, Lei Xu, Xiaoxiao Zheng, Meina Fu, Shuxia Yao, Carlo V. Cannistraci, Keith M. Kendrick, Benjamin Becker

**Affiliations:** Clinical Hospital of Chengdu Brain Science Institute, MOE Key Laboratory for Neuroinformation, University of Electronic Science and Technology of China, Chengdu, China; Biomedical Cybernetics Group, Biotechnology Center (BIOTEC), Center for Molecular and Cellular Bioengineering (CMCB), Center for Systems Biology Dresden (CSBD), Cluster of Excellence Physics of Life (PoL), Department of Physics, Technische Universität Dresden, Dresden, Germany; Center for Complex Network Intelligence(CCNI), Tsinghua Laboratory of Brain and Intelligence (THBI), Department of Bioengineering, Tsinghua University, Beijing, China

**Keywords:** emotional expression, encoding, face recognition, fMRI, long-term memory, multivariate, PCA, vmPFC/OFC

## Abstract

Studies demonstrated that faces with threatening emotional expressions are better remembered than non-threatening faces. However, whether this memory advantage persists over years and which neural systems underlie such an effect remains unknown. Here, we investigated recognition of incidentally encoded faces with angry, fearful, happy, sad and neutral expressions over >1.5 years (*N*= 89). Univariate analyses showed that threatening faces (angry, fearful) were better recognized than happy and neutral faces after >1.5 years, and that the threat-related memory enhancement was driven by forgetting of non-threatening faces. Multivariate principal component analysis (PCA) confirmed the discriminative performance between threatening and non-threatening faces. With an innovative Behavioral Pattern Similarity Analysis (BPSA) approach and functional magnetic resonance imaging (fMRI) acquisition during encoding, we further found that the long-term memory advantage for threatening faces were underpinned by differential neural encoding in the left inferior occipital gyrus (IOG) and right ventromedial prefrontal/orbitofrontal cortex (vmPFC/OFC). Our study provides the first evidence that threatening facial expressions lead to persistent face recognition over periods of >1.5 years and differential encoding-related activity in the visual cortex and medial prefrontal cortex may underlie this effect.

## 1. Introduction

For social species the recognition of previously encountered conspecifics is vital for survival and successful interaction. In humans, faces are presumably the most important stimuli for subsequent recognition. Given the high evolutionary significance of these stimuli, cortical networks specialized in perceiving and recognizing faces develop already during early infancy (Powell, Kosakowski, & Saxe, 2018; Cohen et al., 2019). Nevertheless, the ability to recognize faces varies greatly in the human population. While some individuals can recognize faces following a single exposure over years, others find it nearly impossible to recognize highly familiar faces (Russell, Duchaine, & Nakayama, 2009; Tardif et al., 2019). In addition to individual differences, several characteristics of the facial stimuli can affect subsequent recognition including emotional expression (Bruce & Young, 1986; Haxby, Hoffman & Gobbini, 2000). From an evolutionary perspective, the emotional expression may transmit important information such that threatening facial expressions (e.g., angry or fearful) can signal danger and thus may relate to harm avoidance in the future (Darwin, 1872; Staugaard, 2010).

In support of this evolutionary hypothesis, some experimental studies have demonstrated a recognition advantage of threatening facial expressions across a variety of delays (Grady, Hongwanishkul, Keightley, Lee, & Hasher, 2007; Jackson, Linden, & Raymond, 2014; Pinabiaux et al., 2013; Stiernströmer, Wolgast, & Johansson, 2016; Thomas, Jackson, & Raymond, 2014; Wang, 2013). For example, previous studies consistently reported that faces with threatening expressions (i.e., angry or fearful) are better remembered compared to non-threatening faces in visual working memory (Jackson et al., 2014; Öhman, Lundqvist, & Esteves, 2001; Thomas et al., 2014). Several studies on short-term memory also found a recognition advantage for threatening faces when the memory was tested immediately (i.e., minutes) after encoding (e.g., Grady et al., 2007; Pinabiaux et al., 2013; Wang, 2013). Following a longer retention interval of 24 hours, the recognition memory was better for fearful relative to neutral faces (Wang, 2013). These findings broadly align with numerous studies indicating enhanced recognition for non-facial emotional stimuli (e.g., scenes or words), particularly high-arousing negative ones, which are more vividly and accurately remembered over retention intervals ranging from minutes to years (reviewed in Bowen, Kark, & Kensinger, 2018; Yonelinas & Ritchey, 2015). On the other hand, several studies failed to observe memory enhancement of threatening faces across a variety of delays (minutes to 2 weeks, Anderson, Yamaguchi, Grabski, & Lacka, 2006; Grady et al., 2007; Satterthwaite et al., 2009; Xiu, Geiger, & Kiaver, 2015). These contradictory results may be accounted for by factors such as the lack of statistical power (i.e., small number of subjects and face stimuli), different methodological approaches, and heterogeneity of sample characteristics such as age and gender which may relate to differences in face memory ability (Grady et al., 1995; Sommer, Hildebrandt, Kunina-Habenicht, Schacht, & Wilhelm, 2013). Moreover, various retention intervals in different studies may also have an effect and emotional face recognition after lengthy retention intervals (i.e., years) as well as the underlying neural basis have not been examined.

Against this background, the first aim of the present study was to investigate the emotional expression effects on face recognition over an extended retention interval (>1.5 years) in a large sample of healthy young adults by capitalizing on a large fMRI sample (Li et al., 2019; Liu et al., 2020; Xu et al., 2020; Zhou et al., 2020). 225 college students underwent incidental encoding of faces with emotional expressions (angry, fearful, happy, sad and neutral, each face image was taken from a different actor) during fMRI acquisition. Twenty minutes after the scanning session, all subjects completed an immediate recognition test in which the previously presented face images (i.e., targets) during scanning were intermixed with a new set of emotional faces (i.e., lures) from a different group of actors. On presentation of each image, participants were instructed to indicate whether the image had been shown in the scanning session. A subsample of subjects (*N* = 102) also participated in another surprise face recognition test after a delay of at least 1.5 years to discriminate between the same targets and a new set of emotional faces. We examined whether face recognition was modulated by facial expressions by means of univariate (ANOVA) and data-driven multivariate (principal component analysis, PCA) approaches. The aim of PCA was to complement the conventional, hypothesis-driven univariate approach and further investigate the robustness of the findings by detecting hidden patterns of the trial-wise behavioral responses of the subjects in an unsupervised manner. Based on the previous findings, we expected augmented recognition of faces with threatening expressions, particularly angry and fearful, compared to non-threatening faces after a retention interval of >1.5 years in the univariate ANOVA and a discriminative memory performance between expression conditions by multivariate PCA.

Furthermore, another critical question that remains to be answered is which neural systems supported the expression-associated memory advantage. Previous fMRI studies have revealed that the long-term memory advantage for emotional stimuli (i.e., scene) recruits memory-related medial temporal lobe (MTL) systems, stimulus-specific perceptual systems such as the visual cortex, limbic systems (amygdala) and prefrontal systems during encoding (Cahill et al., 1996; Dolcos, Denkova, & Dolcos, 2012; Dolcos, Labar, & Cabeza, 2005; Erk, Kalckreuth, & Walter, 2010; Ritchey, Dolcos, & Cabeza, 2008). However, few studies have investigated the neural mechanisms of the long-term memory advantage for emotional faces. Given that the ability in general face recognition (Tardif et al., 2019; Wang, Li, Fang, Tian, & Liu, 2012) and emotional expression processing (Calder & Young, 2005; Le Grand et al., 2006) varies considerably in the population, and some people may even use a qualitatively different mechanism to process face stimuli (Tian et al., 2020), it is conceivable that individual differences exist with respect to the long-term emotional face memory and the corresponding neural correlates. Thus, the second aim of the present study was to examine whether brain activation during incidental encoding may associate with the emotional expression effects on long-term face recognition after 1.5 years.

To this end, we initially devised an innovative approach termed Behavioral Pattern Similarity Analysis (BPSA) by separating the subjects into two groups to facilitate the fMRI analyses. In the present study, the BPSA approach measured the similarity of multi-item confidence rating pattern to emotional faces of each subject and the principal component score pattern derived from the previous PCA. This method allowed us to distinguish subjects with or without discriminative face representation of multivariate data based on the significance of similarity so as to inform the subsequent fMRI analyses with a high sensitivity and sufficient power to determine the neural correlates. Next the fMRI analyses were performed by comparing the neural response at encoding between the group of subjects with discriminative patterns and the group of subjects with non-discriminative memory patterns to uncover the specific regions sensitive to the long-term memory advantage of threatening faces. Since both emotional enhancement of memory and emotional face processing have been strongly linked to limbic and prefrontal systems, specifically regions engaged in emotional reactivity and value processing such as amygdala and orbitofrontal cortex (OFC) (Hariri, Mattay, Tessitore, Fera, & Weinberger, 2003; Kark & Kensinger, 2019; Kensinger & Schacter, 2008; Rolls, 2019; Vuilleumier, Richardson, Armony, Driver, & Dolan, 2004), we hypothesized that encoding-related activity in these systems may underlie individual differences in the long-term effects of emotional expressions on face recognition.

## 2. Materials and Methods

### 2.1 Subjects

A total of 102 (53 males, age range: 20-32) healthy, young right-handed Chinese students participated in this study which was part of a large-scale fMRI project (e.g., Li et al., 2019; Liu et al., 2020; Xu et al., 2020; Zhou et al., 2020). Due to incomplete behavioral and fMRI data (*N* = 7), extremely low hits and false alarms (hits < 1 and false alarms < 1, *N* = 4), or excessive head motion during fMRI scanning (*N* = 2), data from 13 subjects were excluded from both behavioral and fMRI analyses, resulting in *N* = 89 subjects (44 males, mean age = 23.80 ± 2.39 years) in the final analyses. Details on recruitment protocols and quality assessments are provide in the **Supporting Information**. The study was approved by the local ethics committee at the University of Electronic Science and Technology of China and in accordance with the latest revision of the Declaration of Helsinki. Written informed consent was obtained from each subject.

### 2.2 Stimuli

A total of 150 face stimuli were selected from two validated Asian facial expression databases: Chinese Facial Affective Picture System (Gong et al., 2011) and Taiwanese Facial Expression Image Database (TFEID) (Chen and Yen, 2007). Facial expressions included angry, fearful, sad, happy and neutral (each from 30 different individual actors, 15 males). All facial stimuli were gray-scaled and covered with an oval mask to remove individual features (e.g., hair). The 150 faces were evenly divided into three sets matched regarding arousal and valence rated by an independent sample (*n* = 20, 10 males, mean age = 21.2 ± 0.70 years) before the experiment. Within each face set, the arousal ratings of emotional faces (angry, fearful, happy, sad) were higher as compared to neutral faces (all *ps* < 0.001), whereas arousal ratings between emotional faces did not differ (all *ps* > 0.05).

### 2.3 Experimental procedure

The present study employed a multiple-stage procedure including an incidental encoding phase and a subsequent memory phase (**Figure.1**). All subjects initially underwent an event-related fMRI paradigm using an emotional face processing task (i.e., incidental encoding) between August, 2016 and October, 2017 (Time 1, T1). Fifty facial stimuli (set 1) were repeatedly presented over two subsequent runs with different pseudorandom sequence, balanced for facial expression and gender (5min 12s per run). Stimuli were shown for 2500ms during which the subjects were required to judge the gender of the face by button press. After each trial, a jittered fixation cross was presented for 2000–5600ms (mean ITI = 3800ms, **see Figure. 1**). Stimuli were presented via E-prime 2.0 (Psychology Software Tools, USA, http://www.pstnet.com/eprime.cfm). Twenty minutes after fMRI acquisition, subjects were asked to complete a surprise recognition memory test (immediate test) outside the scanner in which the 50 previously presented faces (set 1, targets) from the fMRI paradigm were intermixed with 50 new faces (set 2, lures). Subjects were instructed to indicate whether each face had been shown during the fMRI acquisition (forced choice: old versus new). Emotional arousal ratings for each face were additionally assessed after the immediate old/new recognition test using a 9-point Likert scale (1 = very weak to 9 = very strong). After a retention interval of >1.5 year (interval range: 653-1113 days), 102 subjects agreed to participate in a surprise recognition test (delayed test) between July, 2019 and August, 2019 (Time 2, T2) in which target faces were intermixed with another set of 50 new faces (set 3, lures). In the delayed test, subjects were asked to rate their recognition confidence on a six-point scale (old vs. new; 1 = definitely new to 6 = definitely old, **see Figure. 1**). The confidence rating approach was employed in the delayed test given that it reflects the strength and quality of the memory more precisely (Aly & Turk-Browne, 2016; Stretch & Wixted, 1998), and thus was more sensitive as compared to the categorical old/new judgement approach. Moreover, this allowed us to conduct multivariate analysis on the delayed test data with increased power. The delayed recognition memory test was carried out online via SurveyCoder 3.0 (https://www.surveycoder.com/).

**Figure. 1.**
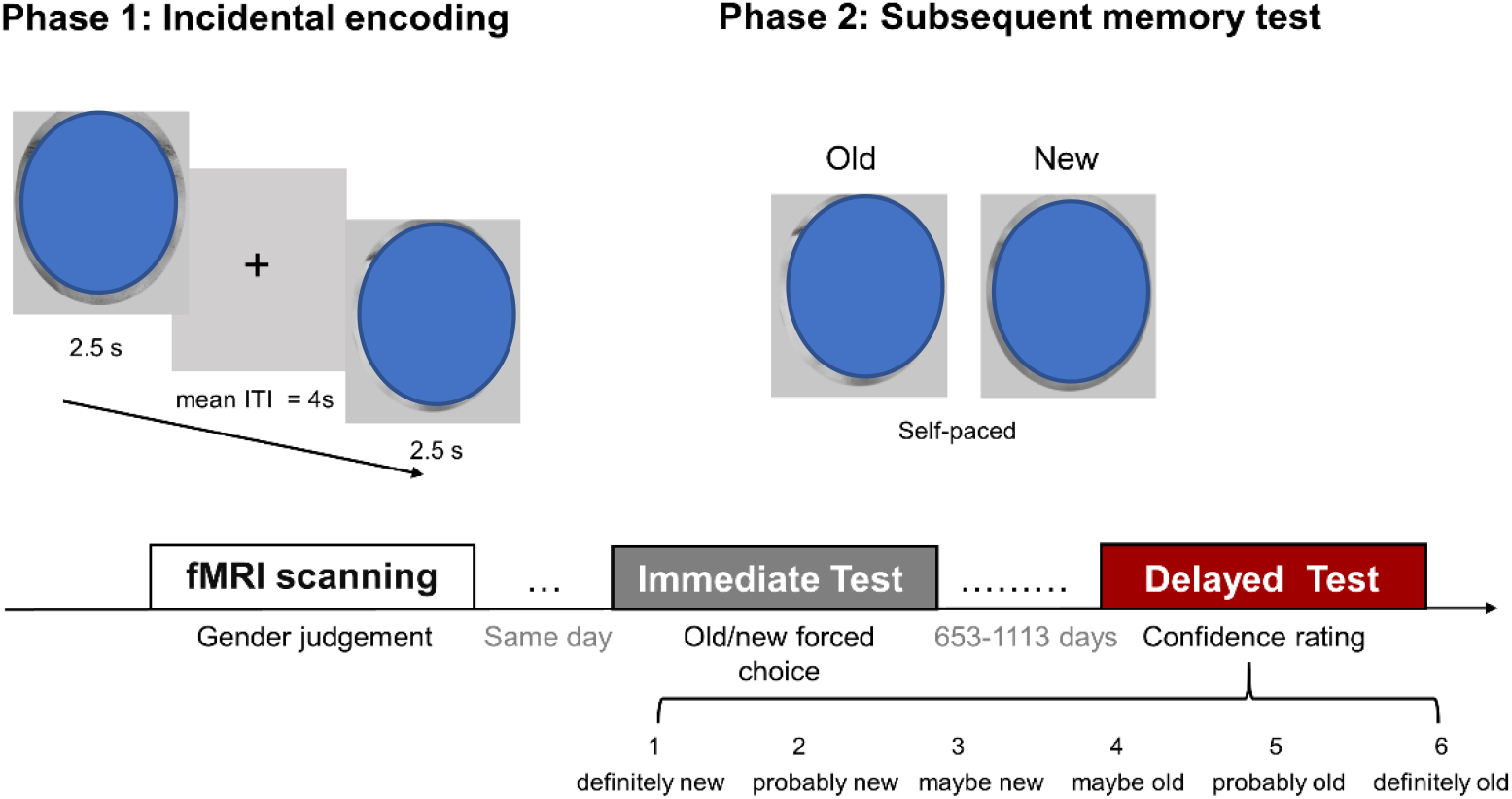
Experimental design and stimuli. Upper panel: Face stimuli in the encoding and subsequent memory stage. Lower panel: Experimental procedure and the behavioral measures. Example face pictures are shown that are not part of the original databases. Consent for publication has been obtained from the individuals presented in the figure.

### 2.4 MRI data Acquisition

MRI data were obtained on a 3T GE MRI system (General Electric, Milwaukee, WI, USA). Functional images were acquired with a gradient echo-planar imaging pulse sequence (39 slices; repetition time (TR) = 2000 ms; echo time (TE) = 30 ms; slice thickness = 3.4 mm; spacing = 0.6 mm; field of view (FOV)= 240 × 240 mm2; flip angle = 90°; matrix size = 64 × 64). Each run of the emotional face processing task consisted of 173 volumes. High-resolution whole-brain T1-weighted images were additionally acquired to improve normalization of the functional images (spoiled gradient echo pulse sequence; 156 slices; TR = 6 ms; TE = 1.964 ms; thickness = 1 mm; FOV = 256 × 256 mm2; flip angle = 9°; matrix size = 256 × 256).

### 2.5 Behavioral data analysis

#### 2.5.1 Univariate approach

To assess whether subjects generally remembered faces immediately after encoding as well as >1.5 years later, a general sensitivity index A-prime (*A’*) was initially computed and compared with chance performance (0.5) (details see **Supporting Information**). Hit rates were subjected to a 2 × 5 repeated-measures ANOVA with the factors time of assessment (immediate vs. delayed) and emotional expression (angry vs. fearful vs. happy vs. neutral vs. sad) to examine the interaction effects between retention interval and facial expression. For both tests, hit rates were defined as the ratio of target faces correctly identified as old. For the delayed test, ratings of 4, 5 and 6 were considered as correctly identified. Post-hoc tests for significant interactions with Holm-Bonferroni correction were conducted to examine the facial expression effects within the immediate or delayed test, and planned two-tailed t tests were performed to determine changes of memory between immediate and delayed recognition in each facial expression condition. Notably, the analyses focused primarily on hit rates because: (*i*) it allows a direct comparison between immediate and delayed test performance, and (*ii*) it focused on the corrected responses of target faces thus establishing a link between the recognition performance and the fMRI encoding process (in which only target faces were included), aligning with the goal of the present study which was to reveal the neural basis during encoding that might contribute to long-term emotional face recognition. We further compared the false alarm rates with chance level as well as among the expression conditions within each test to control for effects of a higher tendency to judge lure items as previously seen. Analyses were computed in SPSS (IBM, SPSS version 20, 2011).

#### 2.5.2 Multivariate approach

Given that the hypothesis-driven univariate analysis (e.g., ANOVA) might dismiss individual differences related to response variability due to averaging, we further investigated the robustness of long-term emotional expression effects using a data-driven unsupervised multivariate approach. The schematic of multivariate analyses is shown in **Figure. 2**. PCA was applied given that it is one of the most widely used unsupervised multivariate machine learning algorithms for exploring the hidden pattern of multidimensional data (Ringnér, 2008), which allowed us to visualize similarities and differences between individual behavioral responses (**Figure. 2A, 2B**). Specifically, the confidence ratings of all target face trials in the delayed test were plotted in a reduced two-dimensional space composed by principal component 1 (PC1) and PC2 (details see **Supporting Information**). PC1 explains the highest variance and represents the most discriminative dimension. We thus expected that trials with the same facial expression (color coded) would dominate separate regions within the reduced space along the PC1 axis, representing a discriminative expression-specific face memory pattern (**Figure. 2B**). To further assess the significance of the segregation, we next employed a recently proposed measure ‘trustworthiness’ which compares the indicator of PC1 segregation (Area Under the ROC Curve, AUC) with a null distribution of AUC values derived from 1000 permutations of shuffled emotion labels (minimum two-tailed *p*-value: 0.05, **Figure. 2B**) (details on this approach see Durán et al., 2021). Finally, Wilcoxon Signed-Rank Tests (two-tailed) with Benjamini-Hochberg-adjusted correction were performed to compare responses between expression conditions. Multivariate analyses including PCA, trustworthiness and Wilcoxon Signed-Rank Tests were implemented in PC-corr MATLAB code (https://github.com/biomedical-cybernetics/PC-corr_net) which has been used in previous studies to successfully discriminate behavioral and omic patterns (Miendlarzewska, Ciucci, Cannistraci, Bavelier, & Schwartz, 2018; Ciucci et al., 2017). The multivariate PCA in combination with the univariate approach thus facilitated a sensitive determination of emotional expression-specific face recognition memory over an extended retention interval. In further exploratory analyses, the PCA was also applied to data from the immediate recognition test to provide a comparison with the delayed test.

**Figure. 2.**
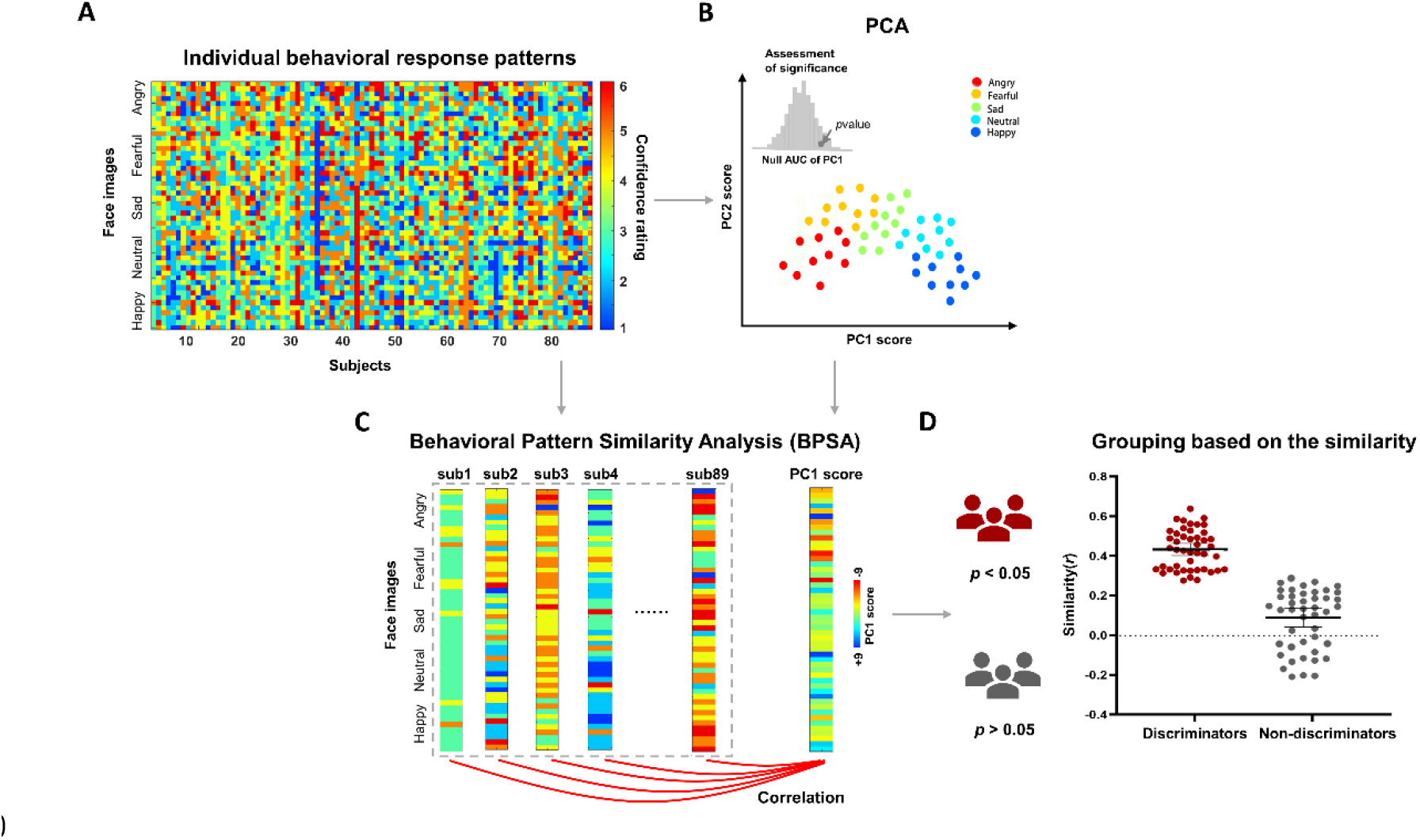
Schematic presentation of the major steps in the multivariate investigation of the long-term emotional face recognition memory. (***A***) Representation of individual behavioral response patterns for all subjects. Each column represents the confidence rating pattern for one subject. Rows from top to bottom indicate face images with expressions angry, fearful, sad, neutral and happy. Color blue to red designates the confidence rating responses from 1 to 6. (***B***) PCA dimensionality reduction and expected discriminative pattern of the facial expression conditions with sample plotted in the 2D reduced space. The inset shows the assessment of significance of segregation using a non-parametric permutation test (i.e., trustworthiness). (***C***) Behavioral Pattern Similarity Analysis (BPSA) between each subject’s response pattern and the PC1 score pattern (template) derived from PCA. The PC1 template was constructed excluding the individual whose correlation was calculated with the template for *N* = 89 times. (***D***) Expected grouping results of subjects based on the BPSA analysis (i.e., one subgroup exhibits discriminative response pattern for facial expression conditions, and one subgroup does not.)

#### 2.5.3 Subject grouping – Behavioral Pattern Similarity Analysis (BPSA)

To account for individual differences in representing a discriminative memory pattern in the delayed test, and especially the lack of differential face memory performance of emotional expressions in some subjects **(Figure. 2A**), we employed a Behavioral Pattern Similarity Analysis (BPSA) approach to characterize the emotional face representation on the individual level. The BPSA is based on multivariate pattern analysis (MVPA) which was proposed for multivariate fMRI data analysis (Haxby et al., 2001; Haxby, 2012) and Inter-Subject Correlation (ISC) analysis which was intended to capture the inter-subject synchronization in brain regions (Hasson, Nir, Levy, Fuhrmann, & Malach, 2004; Hasson et al., 2009; Hasson et al., 2010). The BPSA integrates previous methods in behavioral data (especially data derived from PCA) and includes the following procedures: (1) select a method for unsupervised dimensionality reduction of the multivariate data (without loss of generality in this study we consider PCA, however we wish to stress that any method can be employed, **Figure. 2B**) and consider a dimension of data representation as response template that proves to offer a discriminative pattern between the samples. In our study, the projections of the samples on the first dimension of PCA embedding (which are the PC1 scores of the PCA on confidence ratings for targets of the delayed test) was extracted as a response template, which represents a synthetic meta-subject associated to the discrimination of the face images by confidence rating (**Figure. 2C**, right), (2) a measure of similarity (without loss of generality in this study the Pearson’s correlation coefficients) between each subject’s confidence rating pattern and the PC1 score template was calculated to evaluate the extent to which the group-level emotional expression-specific face representation manifested at the individual level (**Figure. 2C**), (3) the subjects who showed significant similarity (*p* < 0.05, according to Pearson correlation) with the PC1 template were considered discriminators exhibiting a discriminative expression-specific face memory effect and the subjects who did not show similarity (*p* > 0.05, according to Pearson correlation) were considered non-discriminators (**Figure. 2D**): This method thus provided us with behaviorally separable groups on facial expression representation which were next used to inform fMRI analyses to uncover the neural mechanisms underlying the long-term emotional expression effect for face recognition. To avoid inflation of the similarity calculation, the PC1 template was constructed employing a leave-one-subject-out approach, and specifically the individual whose correlation was calculated was excluded from constructing the template.

### 2.6 fMRI data analysis

#### 2.6.1 Image preprocessing

The functional MRI data was preprocessed and analyzed using SPM12 (Statistical Parametric Mapping, https://www.fil.ion.ucl.ac.uk/spm/software/spm12/). The first ten volumes were discarded to allow for MR equilibration. The remaining functional images were realigned to correct for head motion, co-registered with the T1-weighted structural images and normalized to Montreal Neurological Institute (MNI) standard template using a two-step procedure including segmentation of the brain structural images and application of the resultant transformation matrix to the functional time-series. The resampled voxel size of functional data was 3 × 3 × 3 mm^3^. Finally, the images were spatially smoothed using a Gaussian kernel with full-width at half-maximum (FWHM) of 8mm.

#### 2.6.2 Statistical analyses

To identify the neural substrates associated with the long-term emotional expression effects on face memory, a whole-brain ANOVA model with emotional expression (e.g., threatening vs. non-threatening faces determined by ANOVA and PCA on the behavior performance) as within-subject factor, and group (discriminators vs. non-discriminators determined by BPSA) as between-subject factor was employed. To this end, the first-level contrast of interest (i.e., threatening vs. non-threatening) was modeled using separate onset regressors for all trials with threatening vs. non-threatening faces, and convolved with the conventional hemodynamic response function (HRF). Six motion parameters were added in the design matrix to control for movement-related artifacts. Next the first-level contrast images were subjected to a two-sample, two-tailed t-test comparing the discriminators and non-discriminators with interval day as a covariate. This analysis allowed us to identify the specific regions sensitive to the long-term emotional memory advantage during encoding while controlling for unspecific processes. Significant interaction effects were further disentangled by extracting the parameter estimates from independent masks from the Brainnetome atlas (http://atlas.brainnetome.org/download.html, Fan et al., 2016) and using bootstrap tests to warrant a high robustness. Group-level analyses were conducted using Statistical nonParametric Mapping toolbox (SnPM13, http://warwick.ac.uk/snpm) with permutation-based inferences (5,000 permutations). Significant clusters in the whole brain were determined using a height threshold of *p* < 0.001 (two-tailed) and an extent threshold of *p* < 0.05 (two-tailed) with cluster-based familywise error (FWE) correction (Eklund, Nichols, & Knutsson, 2016; Slotnick et al., 2017).

Given that the amygdala has been strongly implicated in emotion processing and emotional memory formation (Blanchard & Blanchard, 1972; Davis, 1992; LaBar & Cabeza, 2006; LeDoux,1995), we examined effects in the amygdala with increased sensitivity using *a priori* region-of-interest (ROI) analysis. Bilateral amygdala masks were created from the Brainnetome atlas (http://atlas.brainnetome.org/download.html, Fan et al., 2016). Small volume correction was performed using FWE correction with a voxel-level threshold of *p* < 0.05.

## 3. Results

Overall, subjects successfully discriminated target faces (all expression conditions) from lure faces in both the immediate test (mean *A*’ = 0.65, *t*_88_ = 15.74, *p* < 0.001, Cohen’s d = 1.67) and the delayed test (mean *A’* = 0.54, *t*_88_ = 3.70, *p* < 0.001, Cohen’s d = 0.4). Paired t-test further indicated that *A*’ in the delayed test was significantly lower than that in the immediate test (*t*_82_ = −9.72, *p* < 0.001, Cohen’s d = 1.03), suggesting that the general recognition performance decreased with time.

### 3.1 Long-term emotional expression effects on face memory

#### 3.1.1 Univariate results

The ANOVA revealed a significant main effect of facial expression (*F*_(4,85)_ = 7.61, *p* < 0.001, *η*^2^ = 0.26) and retention interval (*F*_(1,88)_ = 27.35, *p* < 0.001, *η^2^* = 0.24) on the hit rates, as well as a significant interaction effect (*F*_(4,85)_ = 4.17, *p* < 0.005, *η*^2^ = 0.16, **Figure. 3A**). Post-hoc tests indicated that facial expression modulated delayed memory recognition (hit rate: *F*_(4,85)_ = 10.18, *p* < 0.001, *η*^2^ = 0.32), but not immediate memory recognition (hit rate: *F*_(4,85)_ = 2.17, *p* = 0.08). Within the delayed recognition, paired t-test further suggested that hit rates for faces with both threatening facial expressions (angry or fearful) were significantly higher as compared to faces with non-threatening expressions (neutral or happy, respectively) (two-tailed *ps* < 0.001, Holm-Bonferroni corrected). In contrast, no significant differences between recognition performance of angry versus fearful as well as neutral versus happy faces were observed, whereas that of sad faces ranged in between (**Supporting Information**). To rule out the possibility that this long-term emotional memory advantage of threatening faces was influenced by variations in the retention interval (ranging from 653-1113 days), repeated-measures ANOVA on hit rate of the delayed test was recomputed including interval day as a covariate and results remained stable (*F*_(4,84)_ = 2.43, *p* = 0.05, *η^2^* = 0.10). Moreover, correlation analyses were conducted between retention interval day and hit rate in each condition and these correlations were not significant (all *ps* > 0.05, **Supporting Information**). To test whether this long-term enhanced memory for threatening faces is a result of higher tendency to respond “old” to threatening faces, two control analyses were conducted. First, we compared the false alarm rates in delayed test with the chance level and the results showed that the false alarm rates for both angry (one-sample t test: *t_88_* = 0.70, *p* = 0.49) and fearful (one-sample t test: *t_88_* = 1.61, *p* = 0.11, for other expression conditions see **Supporting Information**) expression conditions were at chance level, whereas the respective hit rates were higher than chance level (angry: one-sample t test, *t_88_* = 3.00, *p* < 0.005; fearful: one-sample t test, *t*_88_ = 2.16, *p* < 0.05; for other expression conditions see **Supporting Information**). Correspondingly, the comparisons between false alarm rates and chance level in immediate test are provided in Supporting Information. Moreover, the false alarm rates between expression conditions within the delayed test were not significantly different after including interval as a covariate (*F*_(4,84)_ = 1.08, *p* = 0.37; although significant difference without controlling for interval days, *F*_(4,85)_ = 9.85, *p* < 0.001, *η*^2^ = 0.32), whereas the false alarms rates during immediate recognition showed an emotion-specific pattern (*F*_(4,85)_ = 22.15, *p* < 0.001, *η*^2^ = 0.51; post-hoc test see **Supporting Information**). Further control analysis was performed on arousal of the face expressions. A hierarchical regression analysis suggested that the interaction of facial expression and arousal did not reveal a significant effect (*Δ*R^2^ = 0.068, *p* = 0.310) on the hit rate. Together, these results suggest that the long-term recognition memory advantage for threatening faces was not influenced by interval days, false alarm rate, and arousal.

**Figure. 3.**
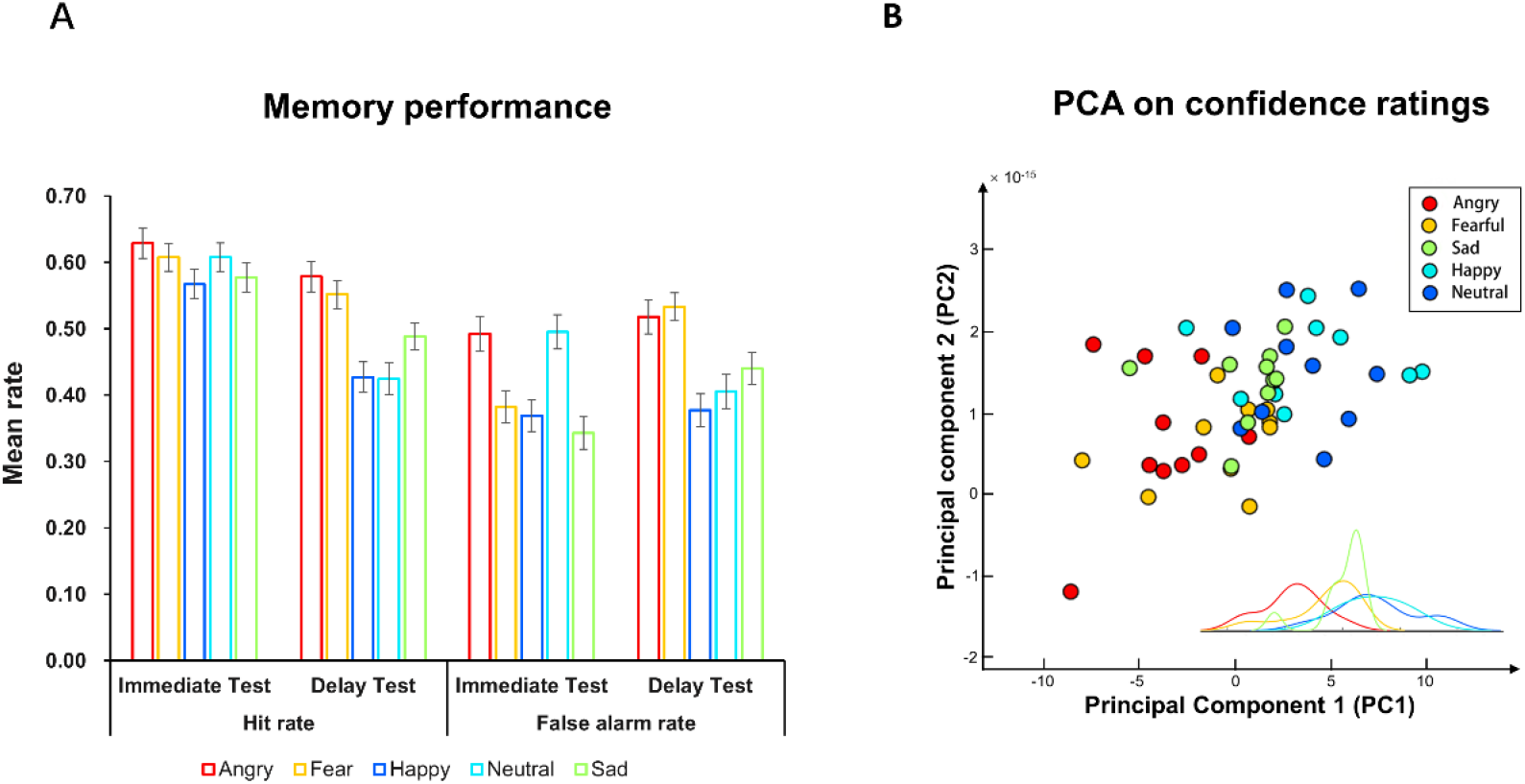
Behavioral results. (***A***) Memory performance for each emotion expression condition in the immediate and delayed memory test as displayed by hit rates and false alarms rates. Error bars depict ±1 SEM. (***B***) PCA on confidence ratings for target faces in the delayed test shows a separation of facial expression conditions (color-coded) on memory performance along PC1 score axis (angry and fearful faces: generally negative scores; happy, neutral, sad faces: generally positive scores). The inset shows the distribution of PC1 scores for each facial expression condition.

To determine changes of memory in each facial expression condition over >1.5 years, pairwise comparisons on hit rates between the immediate and delay test for each facial expression condition were conducted. Recognition performance for happy, neutral and sad faces (two-tailed paired t-test: happy: *t*_88_ = −5.25, *p* < 0.001, Cohen’s d = 0.56; neutral: *t*_88_ = −6.10, *p* < 0.001, Cohen’s d = 0.65; sad: *t*_88_ = −3.21, *p* < 0.01, Cohen’s d = 0.34, **Figure. 3A**) significantly declined during the 1.5-year retention interval, whereas recognition for angry (*t*_88_ = −1.65, *p* = 0.104) and fearful (*t*_88_ = −2.04, *p* = 0.09) faces remained unchanged after Holm-Bonferroni correction.

Together, the results indicated a long-term face recognition advantage of threatening expressions (i.e., angry and fearful) and this advantage was driven by decreased recognition of faces with non-threatening expressions including happy, sad and neutral following a retention interval of 1.5 years.

#### 3.1.2 Multivariate results

Initial inspection of the response patterns (color-coded trial-wise response with blue to yellow as the confidence increased, see **Figure. 2A**) revealed strong individual variations in confidence ratings for expression-specific face images in the delayed test. PCA was then applied to map the confidence ratings of all 50 target face images in the 2D geometrical space of PC1 and PC2. As expected, PC1 had a discriminative variability that accounted for facial expression conditions with polarity: angry and fearful faces (generally negative scores) versus happy, neutral, sad faces (generally positive scores) (**Figure. 3B**). In particular, considering the localization of each face trial along the PC1 axis as visual reference, angry and fearful were separated from neutral and happy while sad was located in between. PC1 explained 12.40% of the variance. The segregation pattern emerging from the amount of variance explained by PC1 was significant compared with the null distribution (trustworthiness/*p* < 0.001). In line with the visual presentation, pairwise non-parametric Wilcoxon Signed-Rank tests revealed significant differences between the facial expression conditions except for angry vs. fearful and neutral vs. happy (see **Supporting Information**). The same PCA procedure was also applied to the immediate test data and failed to detect a discriminative memory pattern along the PC1 axis (see **Supporting Information**, Figure S1). The lack of separation according to facial expression in the PCA is in line with findings from the conventional univariate non-parametric approach (Wilcoxon Signed-Rank tests) which did not reveal significant interaction effects between emotion and recognition performance for the immediate test (**Supporting Information**).

To summarize, our univariate analyses suggested a long-term memory advantage of threatening (i.e., angry and fearful) versus non-threatening (i.e., particularly happy and neutral faces and to a lesser extent sad) faces, and this emotional expression effect was further supported by the results from the multivariate approach.

### 3.2 Distinct long-term emotional face representation between subgroups

Given that the PC1 scores derived from the previous PCA revealed a group-level emotional expression dimension with polarity (angry/fearful: generally negative scores; happy/neutral/sad faces: generally positive scores, see **Figure. 3B**), we next conducted the BPSA by correlating the individual confidence rating pattern with the PC1 score pattern (i.e., template) in which a significant similarity reflects differential memory between threatening versus non-threatening faces on the individual level, whereas a lack of similarity reflects no differential memory between emotional expression conditions. The BPSA approach successfully separated the subjects into discriminators (*N* = 43, 21 males, *rs* > 0.28, *ps* < 0.05) and non-discriminators (*N* = 46, 23 males, *rs* < 0.28, *ps* > 0.05). To further confirm the separable emotional face representations between the two subgroups, we calculated the similarity between each discriminator’s multi-item discriminability pattern and the mean pattern of the discriminators (within-group similarity) and compared it with the similarity between each non-discriminator’s multi-item discriminability pattern and the mean pattern of the discriminators (between-group similarity). As expected, the within-discriminator group similarity was significantly higher than between-group similarity (*t*_87_ = 14.20, *p* < 0.001, Cohen’s d = 3.01), suggesting that the non-discriminators’ discriminative patterns significantly deviated from that of the discriminator group. The distinct memory pattern between the two groups was further confirmed using a univariate method by comparing the confidence ratings of the angry/fearful (bin), sad and happy/neutral (bin) faces, given that both univariate and multivariate PCA results suggested a separation of recognition performance among the three conditions. A significant 2-way interaction was detected (*F*_(2,86)_ = 20.49, *p* < 0.001, *η*^2^ = 0.19, **Figure. 4A**). Post-hoc tests showed significant differences between all pairs of the expression conditions in the discriminators, but no differences in the non-discriminators (**Figure. 4A, Supporting Information**). The discriminators and non-discriminators did not differ with respect to gender, age, retention interval, arousal rating for each facial expression condition and overall recognition performance, arguing against confounding effects of these variables on the long-term memory advantage of threatening expressions (for detailed results, see **Supporting Information**). The distinguishable response patterns of the two subgroups thus allowed us to conduct the fMRI analysis with sufficient power by group comparison.

**Figure. 4.**
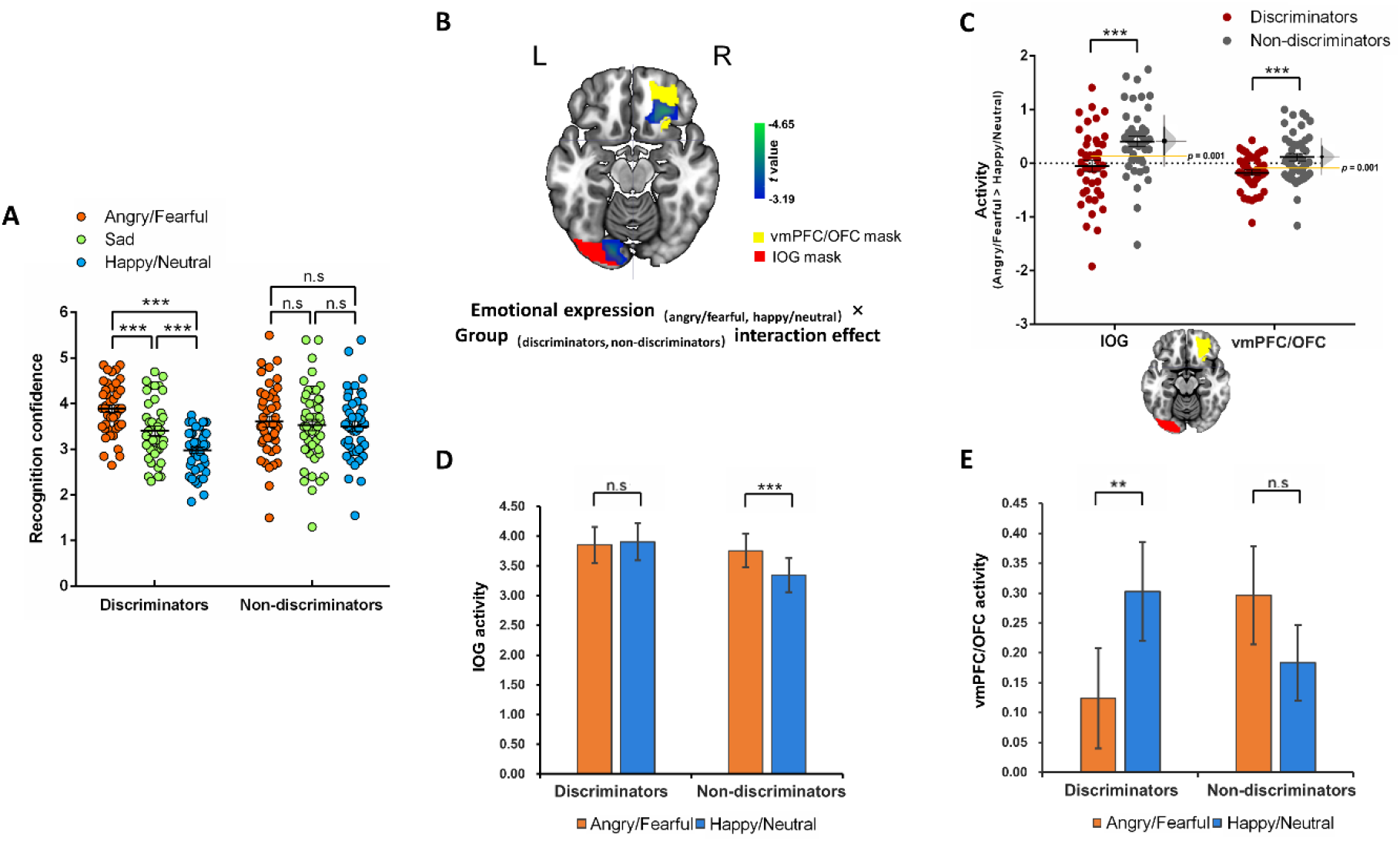
The interaction effect of neural responses at encoding. (***A***) ISC successfully separated the subjects into discriminators (significant differences between angry/fearful, sad and happy/neutral conditions) and non-discriminators (non-significant differences). (***B***) Whole-brain analysis revealed significant emotional expression (angry/fearful vs. happy/neutral) by group (discriminators vs. non-discriminators) interaction effects in the left inferior occipital gyrus (IOG) and right ventromedial prefrontal/orbitofrontal cortex (vmPFC/OFC). The activation map is displayed at *p* < 0.05 cluster-level FWE correction with a cluster-forming threshold *p* < 0.001, overlaid on the masks from Brainnetome atlas which were subsequently used to extract parameter estimates. (***C-E***) Post-hoc tests on the extracted parameter estimates from the left IOG and right vmPFC/OFC (beta estimates) as defined by independent masks from the Brainnetome atlas. Discriminators responded with lower activation for threatening vs. non-threatening faces than non-discriminators in both the left IOG and the right vmPFC/OFC (*C*). The left IOG and the right vmPFC/OFC exhibited distinct neural reactivity patterns such that there was higher reactivity towards angry/fearful faces in the left IOG of non-discriminators (*D*), whereas the discriminators responded less to angry/fearful faces in the vmPFC/OFC (*E*). *** *p* < 0.001, ** *p* < 0.01, Holm-Bonferroni corrected.

### 3.3 Neural basis during encoding

Behavioral results revealed that long-term recognition for angry and fearful faces could be robustly separated from happy and neutral faces across univariate and multivariate analyses, yet no clear pattern emerged for sad faces. Hence, the first-level contrast of interest was modeled as threatening (angry + fearful) > non-threatening (happy + neutral) expressions. The whole-brain group-comparison (discriminators vs. non-discriminators) analysis revealed significant interaction effects in the left inferior occipital gyrus (IOG, *k* = 127; peak MNI coordinates: −18, −82, −19; *t* = −4.65; two-tailed *p*_cluster-FWE_ < 0.05, whole-brain corrected, **Figure. 4B**) and the right vmPFC/OFC (*k* = 131; peak MNI coordinates: 27, 32, −16; *t* = −3.90; two-tailed *p*_cluster-FWE_ < 0.05, whole-brain corrected, **Figure. 4B**). Post-hoc analyses were performed with independent masks that correspond to the location of the IOG (atlas label 205, cluster size = 271, overlapping voxels = 42; **Figure. 4B**) and vmPFC/OFC (atlas label 46, cluster size = 356, overlapping voxels = 64; **Figure. 4B**) from the Brainnetome atlas (Fan et al., 2016). Extraction of beta estimates from the masks showed that the discriminators responded with lower activation for threatening vs. non-threatening faces than non-discriminators in both the left IOG (post-hoc *p* < 0.001, two-tailed Bootstrap test, **Figure. 4C**) and the right vmPFC/OFC (post-hoc *p* < 0.001, two-tailed Bootstrap test, **Figure. 4C**). A further extraction of the estimates of the threatening and non-threatening face conditions respectively from the masks suggested a distinct pattern of neural responses in the left IOG and right vmPFC/OFC. That is, in the left IOG, the non-discriminators showed higher reactivity towards threatening faces compared to non-threatening faces (*t*_45_ = 4.50, *p* < 0.001, Holm-Bonferroni corrected, Cohen’s d = 0.66) while there was no difference in discriminators (*t*_42_ = −0.49, *p* = 0.63, **Figure. 4D**). Conversely, the discriminators responded less to threatening faces compared to non-threatening faces (*t*_42_ = −3.66, *p* < 0.01, Holm-Bonferroni corrected, Cohen’s d = 0.56) whereas the non-discriminators showed no difference in the right vmPFC/OFC (*t*_45_ = 1.72, *p* = 0.09, **Figure. 4E**). These results further suggest that the separation based on the behavioral response patterns allowed the determination of groups with differential neural activity patterns. Against our expectations, the examination of the bilateral amygdala with SVC did not reveal a significant interaction effect.

Additionally, examining the main effects of the emotional expression and group revealed that the bilateral middle temporal gyrus (right MTG: *k* = 295, peak MNI coordinates: 60, −55, −1, *t* = 5.52, *p*_cluster-FWE_ < 0.05, whole-brain corrected; left MTG: *k* = 189, peak MNI coordinates: −54, −58, 2, *t* = 4.50, *p*_cluster-FWE_ < 0.05, whole-brain corrected, see **Supporting Information, Figure S2**) and the left fusiform gyrus (*k* = 130, peak MNI coordinates: −33 −73, −16, *t* = 4.29, *p*_cluster-FWE_ < 0.05, whole-brain corrected, see **Supporting Information, Figure S2**) exhibited higher reactivity towards threatening vs. non-threatening faces during encoding irrespective of group, whereas no significant main effect of group was observed, arguing against unspecific face encoding differences between the groups.

In brief, these results demonstrated that differential reactivity to threatening vs, non-threatening faces in the left IOG and the right vmPFC/OFC during encoding might contribute to the maintenance of long-term memory for threatening faces.

## 4. Discussion

The present study systematically examined the impact of emotional facial expressions during initial encounters on recognition following a retention interval of > 1.5 years. In line with our hypothesis, we found evidence that individuals better recognized threatening faces, particularly angry and fearful ones, as compared to non-threatening faces following the long-term retention interval. The emotional expression-specific recognition advantage was not present directly after encoding and the long-term advantage of threatening faces was driven by decreased recognition of non-threatening faces over the retention period. Multivariate analyses further supported this finding by showing a separation according to emotional face expression following the long-term retention interval but not during immediate recognition. Moreover, the expression-specific face recognition pattern exhibited considerable inter-subject variation and a data-driven ISC demonstrated that approximately half of the subjects demonstrated discriminative emotional face representation (discriminators) on the behavioral level while the other half did not (non-discriminators). Examination of neural activation differences during encoding between these groups revealed that discriminators and non-discriminators exhibited different activation patterns in the IOG and the vmPFC/mOFC in response to threatening vs. non-threatening faces suggesting that different encoding of the face emotions in these regions may precede the differential memory patterns 1.5 years later. Together, the present findings demonstrate that threatening expressions during incidental encounters may facilitate long-term face recognition and that differential encoding in the IOG and vmPFC/mOFC may contribute to expression-associated recognition differences.

Previous studies examining effects of facial expression on face recognition memory used relatively short retention intervals ranging from minutes to weeks (e.g., Anderson et al., 2006; Grady et al., 2007; Pinabiaux et al., 2013; Wang, 2013; Xiu et al., 2015). The present study demonstrated for the first time enhanced memory for faces with threatening expressions relative to non-threatening faces after an extensive period of at least 1.5 years consistent with findings in long-term emotional face recognition after a 24-h delay (Wang, 2013). The long-term memory-enhancing effect of threatening facial expressions is moreover in line with studies showing a memory advantage of negative non-facial (i.e., scenes) stimuli after a retention interval of 1 year (Dolcos et al., 2005; Erk et al., 2010; Gavazzeni et al., 2012). No differences between facial expression conditions were observed in the immediate test, which is consistent with some previous studies showing a lack of emotional facial expression modulation during immediate memory (Anderson et al., 2006; Grady et al., 2007; Satterthwaite et al., 2009; Xiu et al., 2015). However, other studies reported better memory for negative faces during immediate face recognition (Grady et al., 2007; Pinabiaux et al., 2013; Wang, 2013). Methodological differences between experiments may account for this discrepancy. For example, the specific emotional expression tested and the number of face stimuli per emotional category varies between studies (Anderson et al., 2006; Xiu et al., 2015; but see Grady et al., 2007; Wang, 2013). Besides, previous studies only tested a small number of subjects (*N* < 50) which may result in a lack of statistical power. Moreover, the mixed findings from previous studies might also attribute to the heterogeneity of sample characteristics such as age and gender which has been reported to be related to general face recognition memory (e.g., Grady et al., 1995; Sommer et al., 2013). For instance, while a study reported enhanced recognition for fearful compared to neutral faces in adolescents (Pinabiaux et al., 2013), the emotional modulation was neither observed in children (Pinabiaux et al., 2013) nor in older adults (Grady et al., 2007). Additionally, different analytical approaches may have contributed to the divergent results such that the previous studies generally conducted univariate analytic approaches which might dismiss individual differences related to response variability due to averaging. Taken together, differences in the experimental design, sample characteristics and analytic approach may have contributed to the divergent results. Our study partly overcame some of the previous limitations by examining emotional expression effect in a large sample of young adults with balanced gender and incorporating both univariate and data-driven multivariate approach analyses.

The time-dependent effects of emotion on recognition found in our study (i.e., enhanced emotional memory after a long-term delay but not immediately after encoding) were in line with prior studies using non-face emotional stimuli such as words or scenes (Sharot & Phelps, 2004; Sharot & Yonelinas, 2008), which indicated enhanced recognition for negative compared to neutral stimuli after a 24-h delay, but not immediately after encoding. Moreover, these previous studies reported that recognition of neutral stimuli decreased over time while recognition of negative stimuli remained the same after a 24-h retention interval (Sharot & Phelps, 2004; Sharot & Yonelinas, 2008). This pattern resembles our present observation on time- and facial emotion-expression dependent changes in recognition memory between the immediate and delayed test. The time-dependent emotion advantage has previously been robustly demonstrated for non-facial stimuli indicating that the enhanced recognition of emotional stimuli emerges after a delay period, suggesting a consolidation-dependent effect (Cahill & McGaugh, 1998; Talmi, 2013; Yonelinas & Ritchey, 2015). Our observation that threatening face memory persisted while non-threatening face memory decreased after a retention interval of 1 year might thus point to a role of emotional expression-specific consolidation of face memory. From an evolutionary perspective, maintaining recognition of threatening faces over long intervals may represent an adaptive and survival-relevant mechanism (Staugaard, 2010), whereas faces with non-threatening expressions are of lower significance for future encounters thus were prone to be forgotten over time (Dunsmoor, Murty, Davachi, & Phelps, 2015). Although emotion may have beneficial effects on immediate recognition, perhaps due to emotion-associated enhanced selective attention during perception or encoding (Feldmann-Wüstefeld, Schmidt-Daffy, & Schubö, 2011; Gable & Harmon-Jones, 2012; Vuilleumier, 2002), the present findings suggest that the memory advantage for threatening faces increases with retention interval reflecting indirect effects on consolidation.

Notably, additional control analyses suggest that the long-term memory-enhancing effect of threatening facial expressions could not completely be explained by factors such as arousal at encoding or an expression-specific tendency to respond items as previously seen. That is, while an expression-specific variation in the false alarm rates was observed at both, immediate and delayed recall, false alarm rates were similar across the emotion expressions when variations in the retention interval were controlled for and false alarm rates for threatening faces were at chance level in the delayed recognition test.

In addition, previous studies investigating emotional memory advantage emphasized the role of arousal (Bradley et al., 1992; Hamann et al., 2001; LaBar & Phelps, 1998; Sharot & Phelps, 2004). Indeed, arousal may bias processing toward salient information that gains processing priority and thus contribute to enhanced consolidation (Mickley Steinmetz, Schmidt, Zucker, & Kensinger, 2012; Ritchey et al., 2008). However, the enhanced memory for threatening faces in our study is unlikely explained by arousal because a hierarchical regression analysis revealed a non-significant interaction of facial expression category by arousal rating on memory performance. Further comparison of arousal ratings for each expression between discriminators and non-discriminators also indicated that arousal was not significantly different in subjects with different emotional face representation patterns (see **Supporting Information**).

An exploratory analysis further examined the brain systems that may promote the long-term beneficial effects of threatening expressions. To this end we capitalized on the PC1 scores derived from the PCA and individual differences in emotional face representation and separated the subjects into discriminators who demonstrated discriminative expression-specific face memory patterns (threatening vs. non-threatening) and non-discriminators who did not using ISC analysis. The groups that were generated based on their behavioral response patterns were subsequently used to inform the fMRI analyses and thus allowed us to examine neural differences between behaviorally separable groups instead of comparing remembered versus non-remembered stimuli on the individual level. Due to the comparably low hit rate, the latter approach would have suffered from a low robustness of the estimation of the neural correlates while the former approach provided a sufficiently powered strategy to explore the neural basis which may underlie the long-term memory advantage for threatening faces. We observed higher activity in response to threatening (i.e., angry/fearful) faces relative to non-threatening (i.e., happy/neutral) faces in the left IOG in non-discriminators, and lower activity for threatening relative to non-threatening faces in the right vmPFC/OFC in discriminators, suggesting that differential encoding-related activity in these regions may underlie individual differences in the long-term memory advantage for threatening faces. The IOG, also referred to as occipital face area (OFA), plays a prominent role in the core face network which has been suggested to support the initial stage of face-specific processes and to provide input to other face-responsive regions (Haxby et al., 2000; Liu, Harris, & Kanwisher, 2002; Pitcher, Walsh, & Duchaine, 2011). Convergent evidence suggests that this region is involved in the processing of different face properties including face identity and expression (Cohen Kadosh, Soskic, Iuculano, Kanai, & Walsh, 2010; Haxby et al., 2000), such that e.g. transcranial magnetic stimulation targeting this region induces decreased accuracy during simultaneous face identity and expression processing (Cohen Kadosh et al., 2010) or patients suffering from acquired prosopagnosia due to OFA damage exhibit deficits in both facial identity and expression recognition (Calder & Young, 2005; Rossion et al., 2003). In contrast to the OFA, the OFC is a part of the extended face network which supports social-emotional aspects of face processing, specifically value-based emotion and reward information (Haxby et al., 2000; Ishai, 2008; O’Doherty, Critchley, Deichmann, & Dolan, 2003). Several neuroimaging studies reported that negative emotional pictures, including angry and fearful faces elicited strong activation in the OFC (Dougherty et al., 1999; Northoff, 2000; Satterthwaite et al., 2009). Moreover, the prefrontal cortex including the OFC contributes to emotional memory and learning (Kumfor, Irish, Hodges, & Piguet, 2013; Rolls, 2019). A previous study showed a subsequent memory effect for emotional faces in the medial frontal cortex including the OFC immediately after encoding (Xiu et al., 2015). Interestingly, this study further reveals that emotional face expression affects the effective connectivity from IOG to OFC and that the strengths of the influence of the IOG over the OFC is negatively correlated with memory performance of faces with negative, neutral and positive expressions. The findings of the present study that the left IOG and the right vmPFC/OFC exhibited distinct emotion-dependent response patterns in non-discriminators and discriminators suggest that the behavioral dissociation in long-term memory formation of emotional faces may result partly from the two different encoding mechanisms in early face processing regions (i.e., IOG) and value processing regions (i.e., OFC). Notably, on the behavioral level no differences were observed during immediate recognition, suggesting that differential activation in these regions may be associated with individual variation during memory consolidation. Further investigations are invited to explore the interplay between regions during encoding or consolidation in predicting individual differences in long-term emotional face memory.

In contrast to our hypothesis, we did not observe differential coding of threatening vs. non-threatening faces in the amygdala between discriminators and non-discriminators. This might be partly explained by previous observations that the amygdala responds equally to positive, negative and neutral faces during encoding (Adolphs, 2010; Ball et al., 2009). Although previous studies have indicated that amygdala activation at encoding predicted greater subsequent memory for emotional compared to neutral scenes after a 1-year delay (Erk et al., 2010; Dolcos et al., 2005), these findings are based on the neural activity of remembered versus forgotten items for emotional versus neutral pictures. Given that the low number of hits for each expression condition after the long retention interval might not support a sufficiently powered trial-wise fMRI analysis (e.g., Becker et al., 2017), we did not conduct such an analysis but rather employed an individual difference approach which compared all trials in threatening vs. non-threatening conditions instead, which might explain the lack of amygdala findings. Moreover, the main effect of emotional expression suggested that threatening expressions elicited stronger activity than non-threatening faces in fusiform gyrus and MTG during encoding, which is consistent with previous meta-analytic findings showing similar results (Sabatinelli et al., 2011; for a review, see Vuilleumier & Pourtois, 2007).

The findings of the present study need to be considered in the context of the strengths and limitations of the study design. First, the application of data-driven multivariate PCA allowed us to detect hidden pattern in the confidence ratings in a hypothesis-free manner. The consistent results with those found in univariate analysis not only show the robustness of the finding about long-term emotional expression effects, but also illustrate the potential of multidimensional data-driven methods for analyzing behavioral responses. Although the first component retained only 12.4% of the original variance which might be due to the noise associated with the long retention interval, the PCA did generate separate clusters (see Figure. 3B) and the PC1 separation was statistically significant according to a non-parametric trustworthiness test, suggesting that this approach successfully identified distinguishable latent groups of facial expressions based on the recognition confidence ratings. More importantly, the extracted scores of PC1, representing a discriminative variability that accounted for the facial expression categories, promoted the BPSA by correlating the PC1 scores with each subject’s confidence rating pattern. This multivariate method allowed us for the first time to examine whether some individuals were distinct from others in long-term emotional face recognition by examining the individual face representation (i.e., multi-item confidence rating pattern) in the context of a group-level emotion-specific discriminative memory performance (indexed by PC1 scores). Compared to traditional univariate methods, multivariate methods (e.g., the proposed BPSA) capitalize on the multivariate distribution of data which captures a wider range of information about all conditions and has higher sensitivity in revealing the individual differences in data representation (Haxby, 2012). Specifically, the present BPSA took the principal component score pattern as a response template which serves to describe the individual response pattern with a specific definition associated to the reduced dimension, whereas previous studies used mean score/activity pattern as a template (e.g., Aly & Turk-Browne, 2016; Tian et al., 2019). Our method could support a broader examination of multidimensional data (e.g., behavioral, gene or fMRI data) comprising of several conditions by calculating the association between individual feature pattern and the pattern template generated by unsupervised dimensional reduction (e.g., PCA, minimum curvilinear embedding, Miendlarzewska et al., 2018).

On the other hand, the small number of face images and the low number of hits for each expression condition did not permit a sufficiently powered trial-wise fMRI analysis comparing remembered versus non-remembered items as in the majority of previous studies examining emotional memory effects (e.g. Becker et al., 2017; Dolcos et al., 2005). Future studies using a larger set of face stimuli of each expression are needed to further explore the subsequent memory effect of emotional expression. Moreover, to examine the long-term memory effect with higher sensitivity, a dimensional confidence rating was used at the long-term retention interval while the immediate recognition test implemented a forced choice response. Although previous studies used a similar approach to convert dimensional confidence ratings into binary responses (Weymar, Löw, & Hamm, 2011; Xiu et al., 2015), the results of direct comparison between immediate and delayed recognition should be interpreted with caution. Furthermore, previous studies investigating the immediate recognition of neutral faces that were previously presented with threatening (angry or fearful) or non-threatening (happy or sad) expressions and found no differences in recognizing neutral faces with threatening versus non-threatening expressions (Satterthwaite et al., 2009), suggesting that the “first impression” created by facial expression did not affect subsequent recognition of face identity. However, whether such a “first impression” has an effect on long-term face identity recognition despite a neutral expression during recognition remains to be addressed in future studies. Notably, our study uncovered the encoding-related neural basis of the emotional expression effects on long-term face memory. However, the memory enhancement of emotion has also been attributed to consolidation and retrieval processes (Dolcos et al., 2005; Ritchey et al., 2008; Schmidt & Saari, 2007; Sharot, Verfaelli, & Yonelinas, 2007). Investigations of neural mechanism during consolidation and retrieval stage may thus provide a more comprehensive understanding of the formation of long-term emotional face memory. Finally, in line with our research goal the discriminative face recognition pattern was determined on the level of emotion categories. Future studies may aim to further disentangle neural activity during item-specific face memory formation within each emotional expression category or across emotions to explore different encoding patterns for face stimuli that are on average highly memorable.

## Conclusions

Our study provides the first evidence for a recognition advantage of threatening faces after a long-term interval of >1.5 years. Exploratory analyses further suggested that individuals who exhibited the memory advantage for threatening faces showed differential encoding of threatening versus non-threating faces in the left IOG and right vmPFC/mOFC as compared to individuals who did not show a discriminative emotional face memory, suggesting that encoding-related activity in the occipital visual cortex and medial prefrontal cortex may play a role in the formation of long-term emotional face memory. These findings extend the theory of long-term emotional memory towards facial stimuli and shed new light on the encoding-related neural basis of preserved memory for faces with threating expressions.

## Acknowledgments

We thank all subjects who participated in this study.

## Data availability statement

Behavioral data are available from the corresponding author upon reasonable request. The code for the multivariate data analysis PC-corr MATLAB code is available via https://github.com/biomedical-cybernetics/PC-corr_net. Statistical images of main analyses can be found at https://neurovault.org/collections/9071/.

## Supporting Information

Additional Supporting Information may be found online in the supporting information tab for this article.

