## Supporting information for "Medial prefrontal activity at encoding determines enhanced recognition of threatening faces after 1.5 years"

**Manuscript title:**

### SUPPLEMENTARY METHODS

#### Participants

We re-contacted $N$ = 225 healthy subjects that had participated in a large-scale fMRI project during which they underwent structural, resting-state (Liu et al., 2020) and task-based fMRI assessments (Li et al., 2018; Xu et al., 2020; Zhou et al., 2020). The task-fMRI test battery included blocked design tasks assessing pain empathy (Li et al., 2018; Xu et al., 2020; Zhou et al., 2020) and affective modulation of response inhibition as well as an event-related emotional face processing paradigm. The initial MRI data was acquired between August, 2016 and October, 2017 (Time1, T1) and subjects were re-contacted between July, 2019 and August, 2019 (Time 2, T2) to participate in a surprise recognition memory test for the faces from the emotional face fMRI paradigm.

A total of 102 subjects (53 males, aged 20 – 32 years) agreed to participate in the surprise recognition memory test at T2. All participants were right-handed with normal or corrected-to-normal vision. Due to incomplete behavioral and fMRI data, data from $N$ = 7 participants were discarded; data from $N$ = 4 participants were excluded from further analyses due to extremely low hits and false alarms (hits < 1 & false alarms < 1); $N$ = 2 participants were excluded due to excessive head motion during fMRI scanning (> 3 mm or 3 degrees). Consequently, data from $N$ = 89 subjects was included in the final analyses (44 males, mean age = 23.80 ± 2.39 years, age range: 20-32).

**Behavioral data analysis**

*Univariate approach*

A-prime (*A'*) represents a bias-free measure that varies between 0.5 and 1.0 with higher scores indicating better discrimination for identifying target faces from lure faces (Macmillan & Creelman, 1991). *A'* was calculated using the following formula: $1/2 + [p(\text{hit}) - p(\text{false alarm})] \times [1 + p(\text{hit}) - p(\text{false alarm})] / \{4 \times p(\text{hit}) \times [1 - p(\text{false alarm})]\}$ (Wang et al., 2012). Specifically, in the delayed memory test, *p*(hit) denoted the correct recognition proportion of target faces for which subjects responded "6 (definitely old)", "5 (probably old)" or "4 (maybe old)", whereas *p*(false alarm) denoted the proportion of foil faces for which subjects incorrectly responded "6 (definitely old)", "5 (probably old)" or "4 (maybe old)". In the immediate memory test, *A'* was calculated in which *p*(hit) denoted the correct recognition proportion of target faces for which subjects reported "old" and *p* (false alarm) denoted the proportion of foil faces for which subjects incorrectly responded "old". *A'* significantly above chance (0.5) confirms that participants can successfully discriminate the target faces from lure faces.

*Multivariate approach*

PCA was performed to capture the confidence rating pattern of facial expression conditions. The PCA permits dimensionality reduction by extracting groups of covariant features across data and compressing them through an orthogonal linear combination in new variables, which is referred to as principal component (PC). The PCA analysis is able to provide a new representation of the dataset according to n (where n is equal to the number of samples when it is smaller than the number of features; and it is equal to the number of features when it is smaller than the number of samples) metafeatures called PC scores. Each PCi (with i = 1 to n) score metafeature is a linear combination of the original features according to which we can order the samples, and each PCi explains a percentage of the variance in the original data that is uncorrelated to all other PCi. In particular, the PC1 explains a larger percentage of variance than that of PC2, the PC2 larger than PC3, and so forth till the last component. Compressed data can be mapped in a reduced two-dimensional (2D) space composed by the first two components in order to visualize and discriminate groups of data that are separated. In our study, we assessed whether the PC1 score represents a synthetic meta-participant which provides the highest variance associated to the discrimination of the face images by confidence rating and unsupervisedly mapped the face trials in the 2D geometrical space to visualize whether the trials associated to different facial expressions (angry, fearful, sad, happy and neutral) occupy different zones of the space. The 89 participants' raw confidence rating scores (i.e., features) of all 50 trials (i.e., samples) were considered together to define a multidimensional dataset. Specifically, we conducted the PCA analysis according to the following procedure:

1. The 50 (sample) × 89 (feature) matrix was mapped by PCA into a new reduced 2D geometrical space that was composed by PC1 and PC2. This new 2D geometrical space represented the segregation of the trials according to the highest amount of cumulative explained variance of the dataset.
2. In order to check whether trials associated to different facial expressions occupied different areas of the geometrical space, we attributed a different

color to each of the five expression conditions. In practice, we expect that clusters of trials with the same color (facial expression) should dominate different geometrical regions of the 2D space.

3. Next, a non-parametric trustworthiness test was conducted to statistically measure whether the pattern emerging from the amount of variance explained by PC1 is generated at random (Durán et al., 2021). The procedure to compute the trustworthiness exploits a resampling technique based on label-reshuffling. To build a null model distribution, the labels are reshuffled uniformly at random on the embedded points of the PC1 whose location is maintained unaltered in the reduced space. For each random reshuffling, a separability measure value (AUC-ROC in our case) is computed. The collection of all these values is used to draw the null model distribution. This distribution is employed to compute the probability (an empirical *p*-value) to get at random a separation equal or higher than the one detected by using the original labels. If this *p*-value is lower than a significant threshold (that for convention is set to 0.05), then we can trust that the pattern emerging from the amount of variance explained by PC1 is unlikely generated at random.
4. Finally, a pairwise two-tailed Wilcoxon Signed-Rank Test with Benjamini-Hochberg-adjusted correction (for multiple hypothesis testing) was performed for PC1, to statistically assess the extent to which the visible segregation between groups of trials corresponds to a significant discrimination between different pairs of conditions (e.g., facial expression categories).

### SUPPLEMENTARY RESULTS

#### Post-hoc tests on the hit rates in the delayed test

Pairwise two-tailed comparisons for the delayed test revealed that the hit rate of angry faces was significantly higher than that of happy ($t_{88} = 5.05, p < 0.001$), neutral ($t_{88} = 4.84, p < 0.001$) and sad ($t_{88} = 3.31, p < 0.005$) faces, whereas there was no difference between angry and fearful faces ($t_{88} = 1.09, p = 0.279$). The hit rate of fearful faces was also significantly higher than that of happy ($t_{88} = 5.43, p < 0.001$), neutral ($t_{88} = 4.97, p < 0.001$) and sad ($t_{88} = 2.79, p < 0.01$) faces. The results remained significant after Holm-Bonferroni correction for multiple comparison testing. The hit rate of happy and neutral faces was not different ($t_{88} = 0.10, p = 0.923$), while the hit rates of happy ($t_{88} = -2.37, p = 0.02$) and neutral faces ($t_{88} = -2.63, p = 0.01$) were significantly lower than that of sad faces. However, the difference between happy and sad faces disappeared after Holm-Bonferroni correction, whereas that between neutral and sad faces remained significant.

#### Correlation analyses between hit rates and interval days in the delayed test

The correlations between interval days and hit rate in each condition were not significant (angry: $r = 0.17, p > 0.05$; fearful: $r = -0.03$, p > 0.05; happy: $r = 0.13, p > 0.05$; neutral: $r = -0.14, p > 0.05$; sad: $r = -0.02, p > 0.05$).

#### One-sample t test examining chance level of hit rates and false alarm rates at immediate and delayed test

In the delayed test, angry ($t_{88} = 3.00$, p < 0.005) and fearful ($t_{88} = 2.16$, p < 0.05) faces were recognized higher than chance level while the hit rates for neutral ($t_{88}$ =-2.89, p = 0.005) and happy ($t_{88}$ =-3.09, p < 0.005) faces were lower than chance level, and the hit rate for sad faces was at chance level ($t_{88}$ = -0.45, p = 0.66). In the immediate test, the hit rates for all expression conditions were higher than chance level (angry: $t_{88}$ =5.56, p < 0.001; fearful: $t_{88}$ =5.07, p < 0.001; happy: $t_{88}$ =3.09, p < 0.005; neutral: $t_{88}$ =4.97, p < 0.001; sad: $t_{88}$ =3.49, p < 0.005). For false alarm rates, while the angry ($t_{88}$ = 0.70, p = 0.49) and fearful ($t_{88}$ = 1.61, p = 0.11) faces in the delayed test were falsely recognized by chance, the false alarm rates for neutral ($t_{88}$ =-3.59, p < 0.005), happy ($t_{88}$ =-4.97, p < 0.001) and sad ($t_{88}$ =-2.43, p < 0.05) were lower than chance level. In the immediate test, the false alarm rates for angry ($t_{88}$ =-0.32, p = 0.75) and neutral ($t_{88}$ =-0.19, p = 0.85) faces were at chance level, whereas those for fearful ($t_{88}$ =-5.47, p < 0.001), happy ($t_{88}$ =-5.68, p < 0.001) and sad ($t_{88}$ =-7.68, p < 0.001) faces were less than chance level.

**Post-hoc tests on the false alarm rates in the immediate test**

Pairwise two-tailed comparisons for the immediate test revealed that the false alarm rate of angry faces was significantly higher than that of fearful ($t_{88}$ = 5.63, $p <$ 0.001), happy ($t_{88}$ = 5.15, $p <$ 0.001) and sad ($t_{88}$ = 7.18, $p <$ 0.005) faces. The false alarm rate of neutral faces was also significantly higher than that of fearful ($t_{88}$ = 4.11, $p <$ 0.001), happy ($t_{88}$ = 5.47, $p <$ 0.001) and sad ($t_{88}$ = 6.33, $p <$ 0.001) faces. By contrast, there was no difference between angry and neutral faces ($t_{88}$ = -0.11, $p$ = 0.92), fearful and happy faces ($t_{88}$ = 0.55, $p$ = 0.58), happy and sad faces ($t_{88}$ = 1.11, $p$ = 0.27). The results remained significant after Holm-Bonferroni correction for multiple comparison testing. Notably, although there was difference on false alarm rates between fearful and sad faces ($t_{88}$ = 2.13, $p$ = 0.036), the difference vanished after Holm-Bonferroni correction.

**Post-hoc pairwise two-tailed Wilcoxon Signed-Rank tests on the confidence ratings in the delayed test**

Non-parametric Wilcoxon Signed-Rank test showed that the confidence ratings for angry and fearful faces were not different (Wilcoxon sign-rank $z$ = -1.33, $p$ = 0.18). The ratings for happy and neutral faces also did not differ (Wilcoxon sign-rank $z$ = -0.11, $p$ = 0.91). On the other hand, there were significant differences of the confidence rating between facial expressions angry vs. happy (Wilcoxon sign-rank $z$ = -5.25, $p < 0.001$), angry vs. neutral (Wilcoxon sign-rank $z$ = -5.05, $p < 0.001$) angry vs. sad (Wilcoxon sign-rank $z$ = -3.82, $p < 0.001$), fear vs. happy (Wilcoxon sign-rank $z$ = -5.20, $p < 0.001$), fear vs. neutral (Wilcoxon sign-rank $z$ = -5.32, $p < 0.001$), fear vs. sad (Wilcoxon sign-rank $z$ = -2.75, $p$ = 0.006), happy vs. sad (Wilcoxon sign-rank $z$ = -2.11, $p$ = 0.035) and neutral vs. sad (Wilcoxon sign-rank $z$ = -2.55, $p$ = 0.011). The results remained stable after Benjamini-Hochberg adjustment for multiple comparison testing.

**Post-hoc pairwise two-tailed Wilcoxon Signed-Rank tests on the old/new forced choice in the immediate test**

Non-parametric Wilcoxon Signed-Rank test showed that the old/new judgment for angry and fearful faces were not different (Wilcoxon sign-rank $z$ = -1.33, $p$ = 0.18). The ratings for happy and neutral faces also did not differ (Wilcoxon sign-rank $z$ = -0.11, $p$ = 0.91). On the other hand, there were significant differences of the confidence rating between facial expressions angry vs. happy (Wilcoxon sign-rank $z = -5.25$, $p < 0.001$), angry vs. neutral (Wilcoxon sign-rank $z = -5.05$, $p < 0.001$) angry vs. sad (Wilcoxon sign-rank $z = -3.82$, $p < 0.001$), fear vs. happy (Wilcoxon sign-rank $z = -5.20$, $p < 0.001$), fear vs. neutral (Wilcoxon sign-rank $z = -5.32$, $p < 0.001$), fear vs. sad (Wilcoxon sign-rank $z = -2.75$, $p = 0.006$), happy vs. sad (Wilcoxon sign-rank $z = -2.11$, $p = 0.035$) and neutral vs. sad (Wilcoxon sign-rank $z = -2.55$, $p = 0.011$). The results remained stable after Benjamini-Hochberg adjustment for multiple comparison testing.

**Distinct long-term face memory patterns between subgroups**

Post-hoc tests showed significant differences between all pairs of the expression conditions in the discriminative group, but no differences in the non-discriminative group. Specifically, the averaged confidence ratings for faces with angry/fearful expressions were higher compared to sad ($t_{42} = 5.54$, corrected $p < 0.001$, Cohen's d = 0.84) and happy/neutral expressions ($t_{42} = 10.32$, corrected $p < 0.001$, Cohen's d = 1.57), and ratings for sad were higher than happy/neutral expressions ($t_{42} = 3.98$, corrected $p < 0.001$, Cohen's d = 0.61) in the discriminative group, whereas there was no difference in the non-discriminative group (angry/fearful vs. sad: $t_{45} = 0.79$, $p = 0.43$; angry/fearful vs. happy/neutral: $t_{45} = 1.48$, $p = 0.15$; sad vs. happy/neutral: $t_{44} = 0.35$, $p = 0.73$). The discriminative and non-discriminative group did not differ with respect to age ($t_{87} = 0.33$, $p = 0.74$), gender distribution ($\chi^2_{(1)} = 0.01$, $p = 0.92$), interval days ($t_{87} = -1.53$, $p = 0.13$), arousal ratings for each expression condition

($F_{(4,84)} = 0.59$, $p = 0.67$) and the overall recognition performance for all faces ($t_{87} = 1.27$, $p = 0.21$), arguing against confounding effects of these variables. Further control analysis revealed that the different memory patterns among angry/fearful, sad and happy/neutral of the two groups in the delayed test was not present at immediate recognition (interaction on hit rates: $F_{(2,86)} = 0.24$, $p = 0.79$).

### SUPPLEMENTARY FIGURES

**Figure S1.** PCA analysis of forced old/new responses in the immediate test. Facial expression conditions were not separated along PC1 axis according to the scatter plot and the distributions. Wilcoxon Signed-Rank tests also detected no significant differences between facial expression conditions (all *ps* > 0.05).

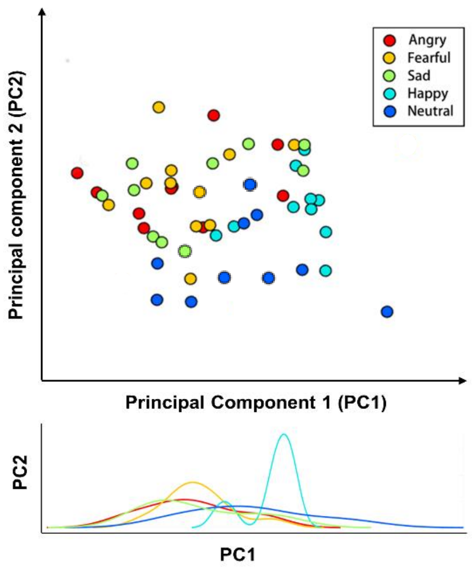

**Figure S2.** Main effect of the emotional expression revealed the bilateral middle temporal gyrus (MTG) and left fusiform gyrus showing higher neural responses to threatening vs. non-threatening faces during encoding. Statistical image is displayed at $p < 0.05$ cluster-level FWE correction with a cluster-forming threshold $p < 0.001$.

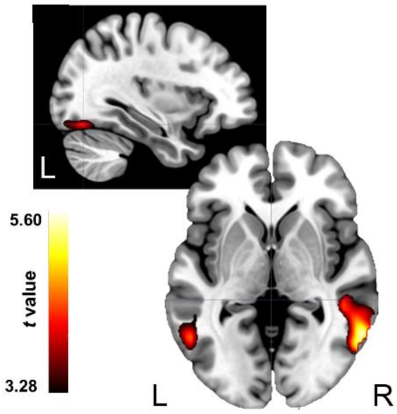
